# Amyloid-β targeting immunisation in aged non-human primate (*Microcebus murinus*)

**DOI:** 10.1101/2022.08.05.502918

**Authors:** Stéphanie G. Trouche, Allal Boutajangout, Ayodeji Asuni, Pascaline Fontés, Einar M. Sigurdsson, Jean-Michel Verdier, Nadine Mestre-Francés

## Abstract

Non-human primates have an important translational value given their close phylogenetic relationship to humans. Studies in these animals remain essential for evaluating efficacy and safety of new therapeutic approaches, particularly in aging primates that display Alzheimer’s disease (AD) -like pathology. With the objective to improve amyloid-β (Aβ) targeting immunotherapy, we investigated the safety and efficacy of an active immunisation with an Aβ derivative, K6Aβ1−30-NH2, in old non-human primates. Thirty-two aged (4-10 year-old) mouse lemurs were enrolled in the study, and received up to four subcutaneous injections of the vaccine in alum adjuvant or adjuvant alone.

Even though antibody titres to Aβ were not high, pathological examination of the mouse lemur brains showed significant reduction in intracellular Aβ without inflammatory or haemorrhagic changes. Moreover, a trend for cognitive improvement was observed in the vaccinated primates, which was probably linked to Aβ clearance. This Aβ derivative vaccine appeared to be safe as a prophylactic measure based on the brain analyses and because it did not appear to have detrimental effects on the general health of these old animals.

## 1. Introduction

The neuropathologies of Alzheimer’s disease (AD) are the most common cause of dementia ^1^. There are no current therapeutic interventions for AD that are clearly disease modifying ^2,3^. A large number of biochemical, histopathological and animal studies as well as numerous genetic, biomarker and clinical reports support the central role of Aβ in the onset of AD pathogenesis. Aβ accumulation in brain seems to be the starting point for the progression of AD, and the most toxic forms of Aβ, soluble oligomers and protofibrils, lead to synaptic damage, neuronal loss and cognitive impairment ^4^.

The amyloid hypothesis has led to the development of passive and active immunotherapy strategies against Aβ, targeting fibrillar aggregates, soluble forms as well as post-translationally modified forms of Aβ. The most simple and attractive of these strategies involved the use of antibodies to slow or prevent disease progression.

Many antibodies have been tested clinically to date and a few of these studies are still ongoing ; however, results over the past two decades have been disappointing ^5^, with the possible recent exception of aducanumab, which is an antibody specific to insoluble fibrillary and oligomeric forms of Aβ ^6^, and Eli Lilly’s anti-pGlu3 Aβ antibody, donanemab ^7^. A few vaccine trials remain active ^8^, but this approach is not without challenges. To avoid autoreactive T cells against Aβ and generate relatively high concentrations of antibodies, it was proposed by various groups, including ours ^9–12^ to develop vaccines by removing T-cell epitopes of full-length Aβ1-42, which may have been responsible for meningoencephalitis in a small subset of AD patients in the original Aβ1-42 vaccine trial ^13^.

The efficacies of Aβ-immunotherapies have been tested in various animal models of AD, resulting in promising data ^14–16^. Most of the immunisation approaches targeting Aβ have been performed in transgenic models that overexpress mutated human amyloid precursor protein resulting in high levels of human Aβ. These mice have normal endogenous levels of mouse Aβ as well. Transgenic mice are ideal for initial screening of potential therapy but prior to clinical trials it is imperative to conduct studies in animals such as non-human primates (NHP) that express normal levels of Aβ, with the same sequence as human Aβ.

Many NHP subspecies develop similar Aβ plaque pathology as seen in AD. Two early findings in four Vervet ^17^ and two Rhesus primates ^18^ have shown that these animals like humans develop an immune response to Aβ 1-40/42 that may have resulted in amyloid clearance ^17^. NHPs are preferred over other animals because their genetics and immune responses are very close to humans.

The ability to test new therapies in a relatively heterogeneous, and thus potentially more generalizable, animal model that recapitulates many aspects of the disease, within a comparatively shorter time, presents an attractive translational opportunity. Mouse lemurs show substantial individual difference in aging. The aged cohorts, similar to those of human, include individuals that age normally, along with animals with age-related cerebral and cognitive pathologies ^19–23^. Our previous study in young animals, comparing vaccination with Aβ 1-42 and various Aβ derivatives showed that all promoted antibody response against Aβ 1-40 and Aβ 1-42 and increased plasmatic Aβ load ^12^. Given that age is a fundamental factor in AD development ^24^, old mouse lemur could offer great insights into the potential translatability of novel and effective therapies. Here, we investigated immunotherapy based on an Aβ derivative, K6Aβ1−30-NH2 administered with alum adjuvant in old mouse lemurs (age between 4 and 10 year-old). Control animals received adjuvant alone. In this small primate (100 g), about 10% of aged animals typically develop Aβ amyloidosis with a large variability in Aβ pathology that is unlikely to be associated with severe cognitive symptoms ^25,26^. The animals in our study were right within the age range when Aβ plaques have started to deposit, and could be considered to be presymptomatic to mildly cognitively impaired, and therefore good candidates for preventive immunisation trial.

## 2. Materials and Methods

### 2.1. Peptides

K6Aβ1−30-NH2 was synthesized at the Keck Foundation at Yale University, as described previously ^9,27,28^. This Aβ derivative maintains the two major immunogenic sites of the Aβ peptide, which are residues 1−11 and 22−28 of Aβ1−42 ^9^. The peptide is amidated on the C-terminus to maintain the immunogenicity of that epitope. The 6 lysyl residues on the N-terminus were added to enhance immunogenicity and further reduce β-sheet content.

### 2.2. Primates

*Microcebus murinus* (mouse lemur primate) is an interesting non-human primate model because it develops Aβ plaques and some hyperphosphorylated tau aggregates with age ^26,29,30^. The animals were born and raised within our breeding facility in Montpellier, France (license approval N°34-05-026-FS). The primates used were disease-free and had never been subjected to any experimental treatment. Thirty-two aged animals were enrolled in the study (age ranging from 4.3 to 9.9 years). The study was conducted according to the guidelines of the European directive 2010/63, and approved by the Ethics Committe Occitanie Méditerranée CEEALR36 (authorization #CE-LR-0713).

### 2.3. Injections and bleeds

Mouse lemurs (n=16) received 100 µg of K6Aβ1-30 subcutaneously in 100 µl alum adjuvant (Adju-Phos, Brenntag Biosector, Denmark). The peptide was mixed with the adjuvant overnight at 4°C to allow it to adsorb onto the aluminium particles. The control group received alum adjuvant alone (n=16) (Table 1).

**Table 1.** Depicts the gender, age as well as the type and number of immunisations received by each of the primates.

| Animal | Sex | Age<br>(years, months) | Group | Number of<br>immunizations |
| --- | --- | --- | --- | --- |
| 129 | Female | 9.9 | Control | 3<sup>1</sup> |
| 174 | Male | 8.9 |  | 2<sup>2</sup> |
| 243 | Male | 7.5 |  | 4 |
| 246 | Female | 7.4 |  | 4 |
| 247 | Female | 7.6 |  | 1<sup>3</sup> |
| 277 | Female | 8.3 |  | 4 |
| 295 | Female | 7.1 |  | 4 |
| 296 | Male | 7.9 |  | 2<sup>4</sup> |
| 313 | Male | 6.6 |  | 4 |
| 314 | Female | 6.9 |  | 4 |
| 325 | Female | 7.7 |  | 4 |
| 327 | Female | 6.9 |  | 4 |
| 332 | Female | 7.1 |  | 4 |
| 333 | Female | 6.7 |  | 3<sup>5</sup> |
| 338 | Female | 7.1 |  | 4 |
| 381 | Male | 4.3 |  | 4 |
| 206 | Male | 7.9 | K6Aβ1-30 | 4 |
| 244 | Female | 7.4 |  | 4 |
| 245 | Female | 7.4 |  | 4 |
| 316 | Male | 7.0 |  | 3<sup>6</sup> |
| 320 | Male | 6.3 |  | 4 |
| 329 | Male | 5.9 |  | 4 |
| 330 | Female | 5.9 |  | 4 |
| 331 | Female | 6.7 |  | 4 |
| 339 | Female | 6.9 |  | 4 |
| 342 | Male | 5.9 |  | 4 |
| 355 | Male | 8.3 |  | 4 |
| 359 | Male | 8.9 |  | 2<sup>7</sup> |
| 362 | Female | 9.5 |  | 3<sup>8</sup> |
| 382 | Female | 4.3 |  | 4 |
| 383 | Female | 4.3 |  | 4 |
| 385 | Male | 4.3 |  | 3<sup>9</sup> |

1 Died 4 months after the 3rd bleed (T2) because of substantial weight loss. The date of death was 6 months after the 1st adjuvant injection.
2 Euthanized 4 days after the 2nd bleed (T1) because of substantial weight loss. The date of death was 25 days after the 1st adjuvant injection.
3 Euthanized 10 months after the 1st bleed (T0) because of substantial weight loss. The date of death was 8 days after the 1st adjuvant injection.
4 Euthanized 18 days after the 2nd bleed (T1) because of substantial weight loss. The date of death was 38 days after the first adjuvant injection.
5 Died 4 months after the 3rd bleed (T2) because of substantial weight loss. The date of death was 6 months after the first adjuvant injection.
6 Euthanized 25 days after the 2nd bleed (T1) because of substantial weight loss. The date of death was 2 months after the first immunisation.
7 Euthanized 8 days after the 2nd bleed (T1) because of substantial weight loss. The date of death was 29 days after the first immunisation.
8 Euthanized 34 days after the 3rd bleed (T2) because of substantial weight loss. The date of death was 3 months after the first immunisation.
9 Died 21 days after the 2nd bleed (T1) because of a deeper subcutaneous injection. The date of death was 42 days after the first immunisation

Alum adjuvant was chosen because it is the most common adjuvant in human vaccines ^31^, and it is considered to be non-toxic at this dose.

The primates received up to four injections. The second, third and fourth injections were administered at 2, 6 and 42 weeks after the first injection (Figure 1). Blood samples were collected prior to vaccination (T0), two weeks after the first injection (T1, week 3) and the third injection (T2, week 7). T3 was at 28 weeks (22 weeks following the third injection) to assess the reversibility of the immune response. T4 and Tfinal (or Tf) were performed at 43 and 44 weeks, respectively. All samples (150-200 µl from the saphein vein) were collected in the afternoon between 2 and 3 pm, when the mouse lemurs are active, to minimize circadian fluctuations. Plasma samples were stored in aliquots at -80°C until further analysis. It was often difficult to obtain sufficient plasma from the lemurs because the blood coagulates very quickly in this species, and it is hard to obtain a large amount of blood from them without causing undue stress.

**Figure 1.**
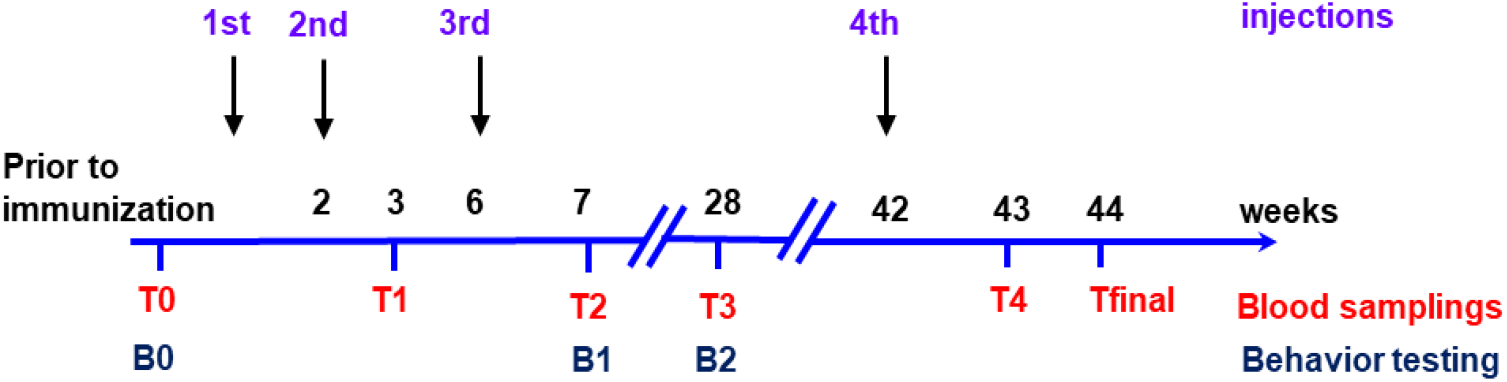
Timeline of the immunisation protocol. The primates (n=16 per group) were bled prior to immunization (T0). Subsequent vaccinations were at 0, 2-, 6- and 42 weeks. Subsequent bleeds were at 3- (T1), 7- (T2), 28- (T3) 43- (T4) and 44 weeks (Tf).

### 2.4 Behavioural testing

The behavioural testing was performed in an experimental room under dim red light, and during the dark phase of the mouse lemur cycle as previously described ^32^. Briefly, working memory was assessed with a three-panel runway maze to measure the working memory ability of the animal. The apparatus includes a departure box and a goal box separated by five compartments. Each choice panel presented three sliding gates. Rear stoppers were used to block the use of two of the three gates in each panel. Mouse lemurs had to perform six consecutive trials per daily session. During each trial, each animal had 10 min to go through all compartments and enter the goal box to get a reward as positive reinforcement. A random sequence of open gates was chosen each day. Only mouse lemurs completing the trial were retested after 2 min for the next trial until a maximum of 6 trials per session. This test was performed at different intervals during the immunisation period (Fig. 1). Animals were tested prior to immunisation (B0), one week after the third injection (B1, 7 weeks) and 22 weeks after the third injection (B2, 28 weeks). Reference errors were analysed to identify an improvement over trials at each behaviour checkpoint.

### 2.5 Antibody titre

To analyse antibody titres, plasma samples were subjected to ELISA as we have described previously ^9,27^. IgM antibodies are usually produced immediately after an exposure to antigens whereas IgG antibodies are associated with a later response Briefly, Aβ1−40 or its derivative K6Aβ1−30 were coated overnight at 4°C onto microtitre wells (0.5 μg/100 μl/well in TBS with 0.1% Tween-20 (TBS-T); (Immulon 2HB, Thermo Electron Corp., Milford, MA). Samples were applied at 1:200 dilution. The signal was detected by an anti-primate IgG or IgM linked to a horseradish peroxidase (1:3000) (Alpha Diagnostics, San Antonio, TX) and tetramethyl benzidine (Pierce Biotechnology, Rockford, IL) was the substrate.

### 2.6 Tissue preparation

At the end of the protocol, the mouse lemurs were killed under deep anaesthesia [ketamine 150 mg/kg) for analysing Alzheimer’s hallmarks and potential inflammatory markers. Blood (Tfinal) was drawn transcardially. Spleens were excised and splenocyte cultures from individual animals were prepared. Brain was removed and bisected sagittally, then the left hemisphere was snap frozen and stored at -80°C until processed for immunological and biochemical studies. The right hemisphere was fixed in 4% paraformaldehyde for 24 hours and then placed in phosphate buffer. Samples were then embedded in paraffin wax. Brain sections were cut on a sliding microtome at a thickness of 6 µm.

### 2.7. Aβ levels in brain and plasma

Extraction of Aβ from brain homogenate was performed as we have described previously in detail ^28^. For measurements of free Aβ1-40 in plasma, 10% dilution of untreated plasma was used, The ELISA procedure was performed as described by the ELISA kit manufacturer (Invitrogen, Camarillo, CA).

### 2.8. Cytokine evaluation

After washing in RPMI 1640 media, lymphocytes were resuspended to a concentration of 5×106 cells/ml in RPMI 1640 media supplemented with 10% foetal bovine serum + Interleukin 2 [20 ng/ml). Cells were plated in 24-well plates and restimulated in triplicate using K6Aβ1-30 peptide (500 µg/ml). At 48 and 72 hours, supernatants were collected and frozen at -20°C until analysed for cytokine production using a multi-analyte ELISArray kit (SABiosciences, Hilden, Germany) with positive and negative controls that allows analysing a panel of 12 human pro-inflammatory cytokines (including IL-4 and IFN-γ). Splenocyte supernatants were added to the plate without dilution. Optical densities at 450 nm were determined spectrophotometrically. Each sample was measured in triplicate.

### 2.9. Histology

Microhaemorrhages were analysed with Perl’s iron stain, which reveals presence of ferric ions. A dark blue staining showed the presence of ferric ions that indicated presence of microhaemorrhages. Perl’s iron stain was performed to detect cerebral bleeding by placing defatted and hydrated sections in a solution containing 5% potassium ferrocyanide and 10% hydrochloric acid for 30 min as we have described previously ^33^. The slides were then rinsed in distilled water, and the sections were dehydrated and coverslipped with Mountex (Histolab, Gothenburg, Sweden).

Brain sections were immunostained with primary antibodies diluted in TBS (Tris Buffered Saline) 1X pH 7.6 that evidenced extracellular Aβ deposits after formic acid pre-treatment. FCA3542 rabbit polyclonal (1/1000, Calbiochem, Darmstadt, Germany) binds to the Aβ1-42 peptide, FCA3340 rabbit polyclonal (1/1000, Calbiochem) binds to the Aβ1-40 peptide, or mouse monoclonal 4G8 (1/1000, Covance, Princeton, NJ) that stained also intracellular Aβ deposits. Congo red staining was performed using a Congo red amyloid stain kit procedure (Sigma Aldrich, St Louis, MO). Pathological tau was evidenced with CP13 (1/50, phospho-serine 202), PHF1 (1/50, phospho-serine 396,404), both generously provided by Peter Davies, and AT100, (1/100, phospho-threonine 212, phospho-serine 214, Innogenetics, Gand, Belgium). Ferritin antibody was used to stain microglia ^34^ (1/500, Sigma) and anti-GFAP (1/100, Dako, Les Ulis, France) for astrocytes, CD3 (1/150) and CD45 (1/250, Abcam, Paris, France) to evidence T cell activation. Appropriate secondary biotinylated antibodies (Clinisciences, Nanterre France) were used followed by avidin peroxidase complex (Vectastain, Vector Laboratories, Les Ulis, France) and 3,3’-diaminobenzidine tetrahydrochloride (DAB) as chromogen for peroxidase activity (Sigma Aldrich). After counterstaining with haematoxylin and dehydration, sections were coverslipped in Mountex (Histolab, Sweden), and analysed with a microscope (Leitz Laborlux S, Wetzlar, Germany).

### 2.10. Image analysis of neuropathological markers

Aβ load was evaluated on sagittal cortex sections by stereological analysis (Sinq-system, Silver Spring, MD). A computerized video microscopy was used for analysis and consisted of Ludl automated XYZ stage movement (with independent linear encoder providing 0.1 µm measurement accuracy in Z) coupled to a Zeiss Axioskop microscope and an Hitachi 3CDD color video camera coupled to a computer running stereologer software.

Neuropathology in brain sections was quantified blindly with the Mercator Pro software (ExploraNova, La Rochelle, France). This software allows quantification of histological sections and can generate maps of counted objects on whole sections such as extracellular cortical Aβ1-42 deposits (every 150 µm, 12 sections/ animal). For intracellular Aβ and ferritin evaluation, the measurement was the number of labelled cells in the random fields (500 × 500 µm for intracellular Aβ and 200 × 200 µm for ferritin) of the different cortical areas (frontal, parietal, occipital and hippocampus). The sections were randomly chosen at 300 µm intervals and five sections were analysed per animal.

### 2.11. Western blotting

Brain tissue was homogenized in a buffer containing 0.1 mM 2-(N-morpholino) ethanosulfonic acid, 0.5 mM MgSO4, 1 mM EGTA;, 2 mM dithiothreitol, pH 6.8, 0.75 mM NaCl, 2 mM phenylmethyl sulfonyl fluoride, Complete mini protease inhibitor mixture (1 tablet in 10 ml of distilled water; Roche Indianapolis, IN) and phosphatase inhibitors (20 mM NaF and 0.5 mM sodium orthovanadate). The homogenate was then centrifuged (20,000xg) for 30 min at 4°C to separate a soluble cytosolic fraction. Standard sarkosyl extraction of the supernatant and subsequent ultracentrifugation revealed limited if any pellet, indicating lack of insoluble tau in these animals regardless of treatment group. Supernatants were heated at 100°C for 5 min and the same amount of protein was electrophoresed on 12% [w/v) polyacrylamide gel. The blots were blocked in 5% non-fat milk with 0.1% Tween-20 in TBS, and incubated with different antibodies overnight, and then washed and incubated at RT for 1 h with peroxidase-conjugated, anti-mouse or anti-rabbit IgG. Subsequently, the bound antibodies (PHF1, CP13, generously provided by Peter Davies), Tau 5 (Thermo-Fischer, Waltham, MA) were detected by ECL (Pierce Biotechnology, Rockford, IL), and imaged with Fuji LAS 4000. Densitometric analysis of immunoblots were performed by Image Quant Software and the levels of pathological tau was normalized relative to total tau protein.

### 2.12. Statistical analysis

The data were expressed as mean + SEM. All the data were plotted and analysed using GraphPad prism 6.0.1 Normality was assessed with the D’Agostino-Pearson test.

Three panel runway task was analysed based on performance in completed trials relative to the first completed session. A repeated measures two-way ANOVA analysis was performed to see an evolution of reference errors in the immunised group in comparison to the control group at the same interval of the immunisation protocol (from B0 to B2).

For tau and Aβ measurements, data that passed normality test, were analysed with an unpaired t-test, and data that failed the normality test were analysed with the Mann–Whitney test. Both tests were two-tailed. For immunological analysis, IgG and IgM levels were analysed by one-way ANOVA or Kruskal-Wallis test when the data failed normality test.

The level of statistical significance for all tests was p <0.05.

## 3. Results

### 3.1 Antibody Response and Plasma Aβ Levels

Using plasma taken at various time points of the experiment, we determined the levels of anti-K6Aβ1-30 and Aβ1-40 antibodies after immunisation of aged mouse lemurs (Fig. 1). Eighty one percent of adjuvant and 93.7% of immunised animals received at least 3 injections (Table 1). Some death occurs during the protocol in both groups but this did not differ significantly between groups. These deaths were mainly in the oldest mouse lemurs. Prior to vaccination, both groups had IgM autoantibodies that recognized the immunogen K6Aβ1-30 and Aβ1-40 (Fig. 2A, B, Fig. 1 suppl, Fig. 2 suppl).

**Figure 2:**
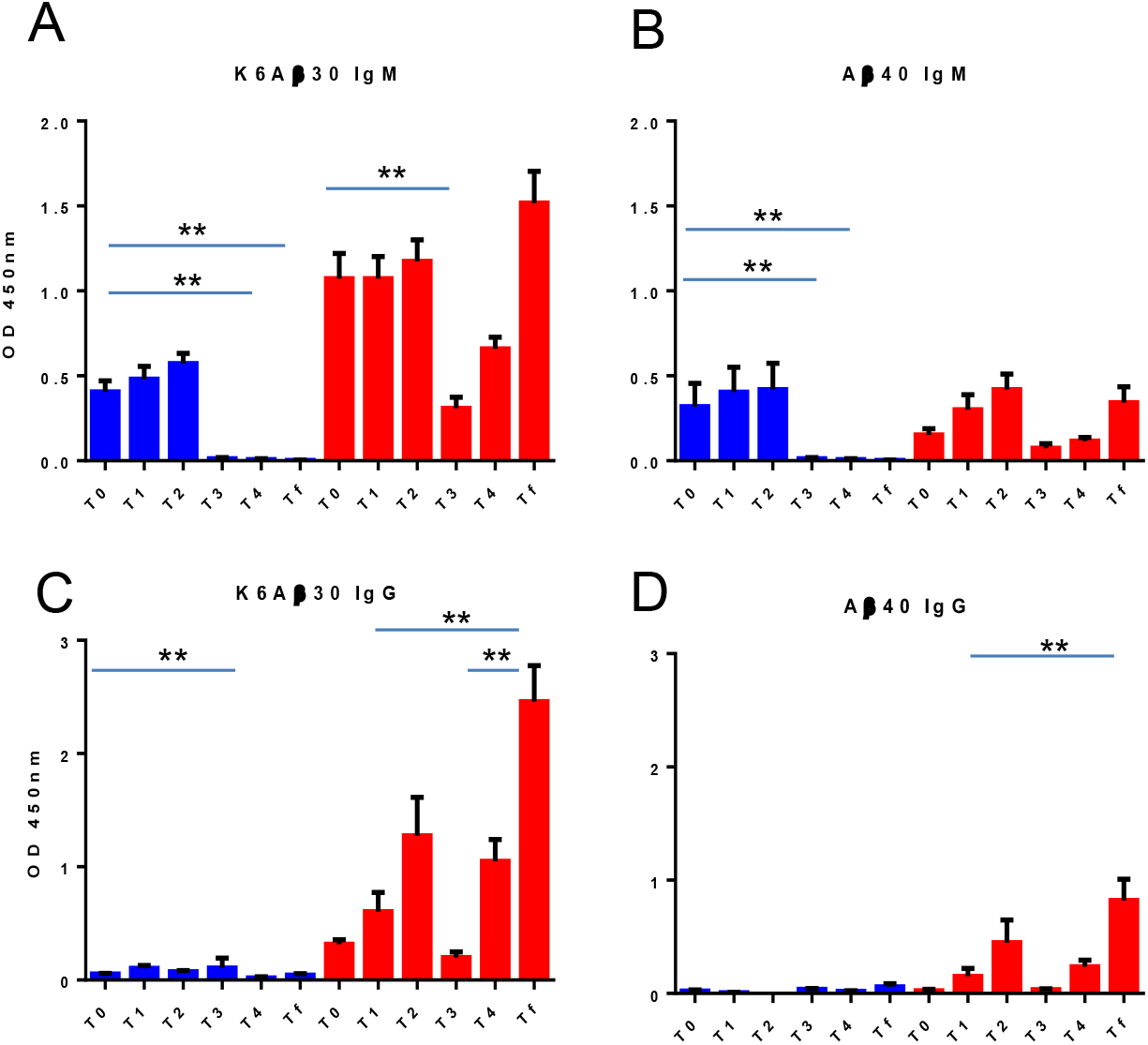
Antibody Response in Aged Primates. (A, B) IgM titre. (C, D) IgG titre. The x-axis depicts which group was tested: adjuvant-treated (blue, n=13), immunised (red, n=14). The y-axis depicts the absorbance at 450 nm. **, p < 0.01

These dissipated over time in the adjuvant group but were detected throughout the vaccination period in the treatment group. The vaccination induced a moderate to strong IgG response against the immunogen and these antibodies recognized Aβ1-40 to a moderate extent (Fig. 2C, D). Levels of both IgM and IgG decreased substantially 22 weeks after the third immunisation (T3) but increased robustly in the two weeks after the fourth immunisation (T4 and Tf). Notably, of the 16 aged animals 44% percent developed Aβ1-40 antibodies at the 6-week time point and 50% after re-immunisation (42-week time point). Four of the 8 responders had high peak antibody levels (Fig. 2 Suppl).

During the protocol, both groups were analysed for Aβ1-40 in plasma at the different time points to determine if the vaccine increased those levels (Fig. 3).

**Figure 3.**
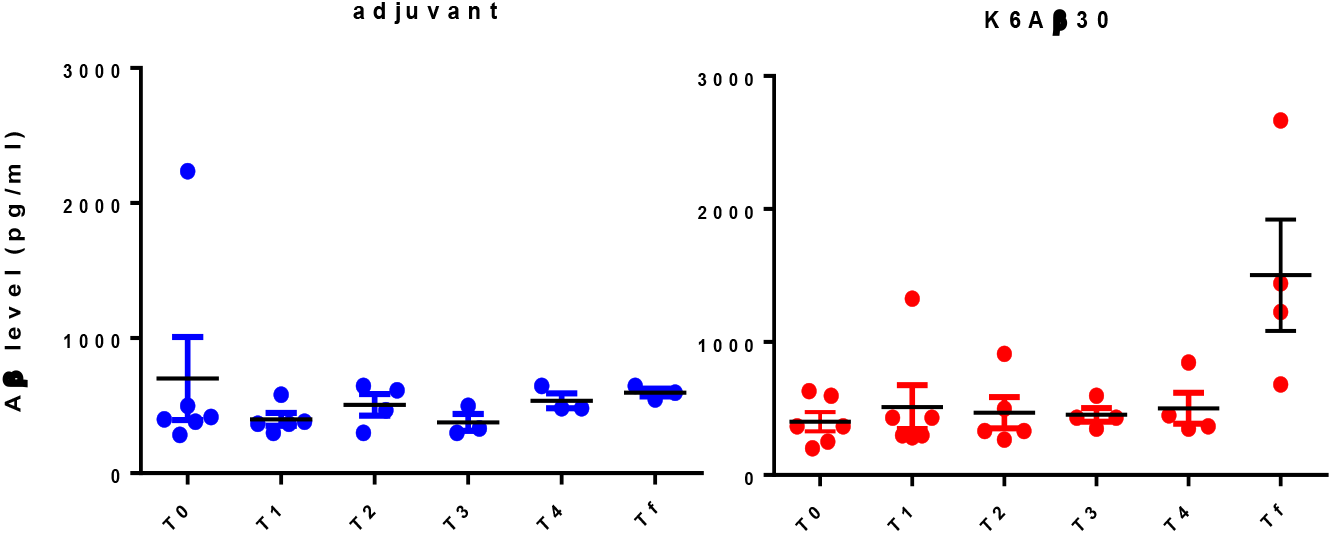
Plasma Aβ1-40 levels in subsequent bleeds from T0 to Tf in adjuvant-treated and immunised animals. The x-axis depicts the timeline of titre check for adjuvant-treated (blue, n=7) and immunised animals (red, n=7). The y-axis depicts the Aβ1-40 level (pg/ml) in plasma. * p <0.05.

Prior to vaccination, both groups, apart from one control animal, had similar Aβ1-40 levels in plasma. During the immunisation protocol, Aβ1-40 levels in the different bleeds did not differ significantly from baseline levels within the groups except for Tf in K6Aβ1−30, and these levels did not differ significantly between the groups. No correlation was observed between plasma levels of Aβ1-40 and antibody response towards Aβ1-40 (data not shown).

### 3.2. K6Aβ1−30 vaccine showed a trend to attenuate cognitive deficits

To evaluate the effect of immunisation on the spatial memory of mouse lemur, we conducted behavioural investigation with the three-panel-runway maze. The number of errors of both groups was evaluated once before and twice during the protocol (Fig. 4A). The adjuvant treated animals showed a tendency to increase the number of errors during this period, suggesting a slight cognitive deficit. During the same period, the K6Aβ1−30 group tended to improve their performance during immunisation (week 7) and returned to baseline after immunisation (week 28).

**Figure 4:**
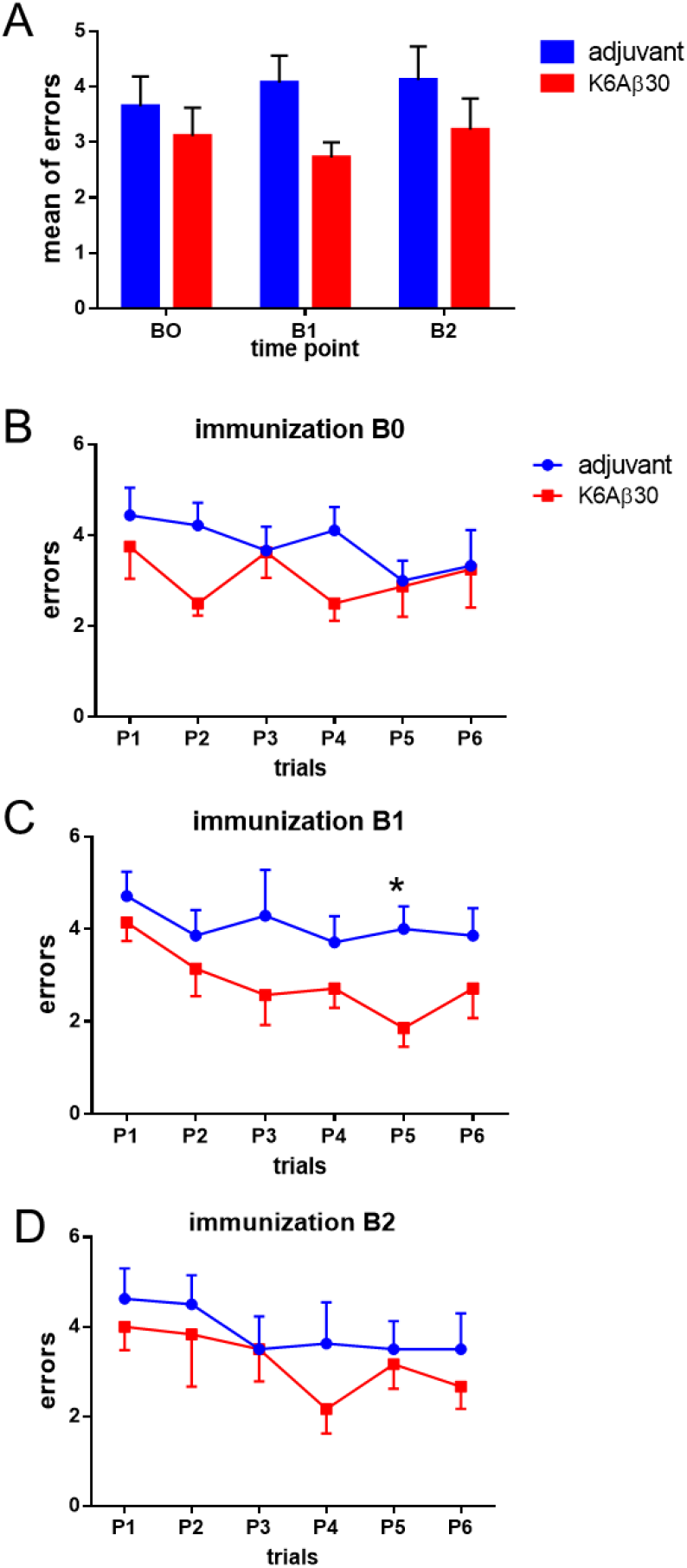
Three panel test evaluation during the immunisation protocol. A -Session to criterion for 3 time points (from B0 to B2). Animals reached the session to criterion (STC) when they complete 5 sessions. Adjuvant group (blue, n=12), immunised group (red, n=12). B-D: reference errors over trials before immunisation B0 (B), during immunisation B1 (C) and after immunisation B2 (D).* p < 0.05.

When we looked at the reference errors over trials, no difference was observed within or between the groups before immunisation (B0, Fig. 4B). After immunisation (B1), immunised animals had a decrease in the number of errors over trials, whereas the control group made more errors than the immunised one (Fig. 4C). Finally, 22 weeks after the third immunisation (B2), immunised and adjuvant-treated groups maintained similar performances (Fig. 4D).

Overall, these results indicate that the K6Aβ1−30 vaccine appeared to have a transient positive effect on spatial memorization.

### 3.3. K6Aβ1−30 immunisation attenuated Aβ deposition

To study the effect of immunisation on brain pathologies in mouse lemur, brains were analysed for Aβ1-40 levels by ELISA. The Aβ levels were variable within each group. No significant difference was observed between the groups (adjuvant: 1.79 + 0.25 ng/ml, immunised 1.73 + 0.25 ng/ml (mean + SEM)). Antibody response towards Aβ1-40 did not correlate with plasma levels of Aβ1-40. Also, peak Aβ antibody response did not correlate with brain Aβ levels as measured by ELISA.

Aβ plaque burden was also assessed by histopathological analysis of the 32 mouse lemurs. As a negative control for Aβ clearance, archived brain tissue from age-matched mouse lemurs was used (n=7). Brain sections were immunolabelled with anti-human Aβ antibodies or labelled with Congo red to assess diffuse or fibrillary Aβ plaque load, respectively. One out of 16 adjuvant-treated animals and one out of 16 immunised ones exhibited amyloid deposits. Percentage of Aβ load evaluated by stereology showed that the immunised animal had an amyloid burden two times less than the adjuvant treated animal, and lesser Aβ load when compared with the 7 age-matched controls (Fig. 5A). Mapping analysis confirm that amyloid plaques were less numerous in the immunised animal than in the adjuvant treated one (p < 0.01, Fig. 5B). The adjuvant control animal had a similar number of Aβ plaques as the age-matched animals.

**Figure 5.**
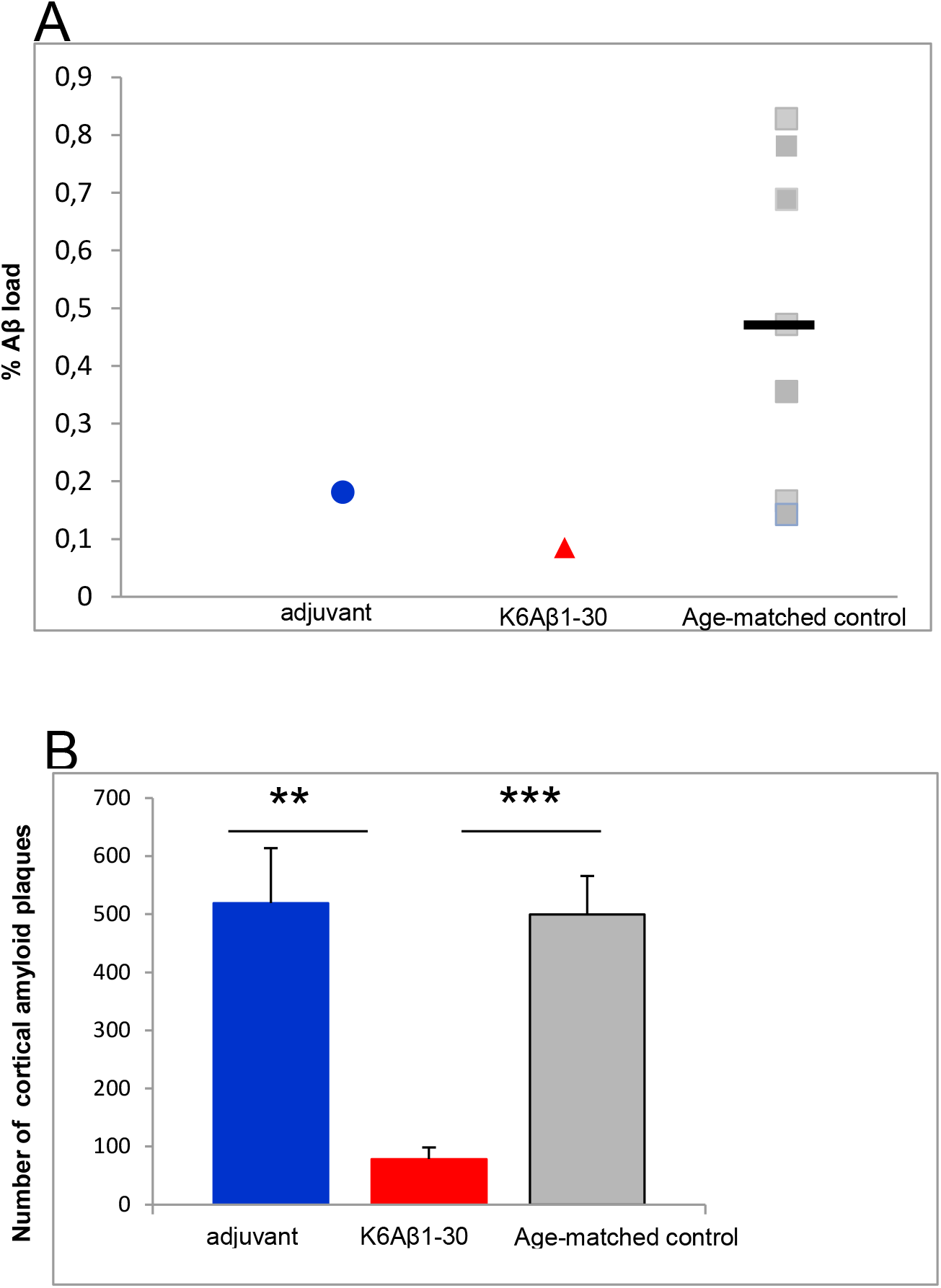
Aβ cortical plaque burden in adjuvant –treated (n=1), immunised (n=1) and aged-matched control (n=7) mouse lemurs. A: percentage of Aβ burden, B: number of Aβ cortical plaques. **, p < 0.01 for adjuvant versus immunised one. *** p < 0.001 for aged lemurs with Aβ plaques versus immunised animal.

A closer examination of the plaques revealed that their features were different. The adjuvant-treated mouse lemur had numerous mild focal plaques (Aβ42 and 4G8 positive) (Fig. 6A, C) with some being Congo red positive (data not shown), whereas the immunised animal had more diffuse deposits (Fig. 6B, D). Compact plaques were not observed in either group.

**Figure 6.**
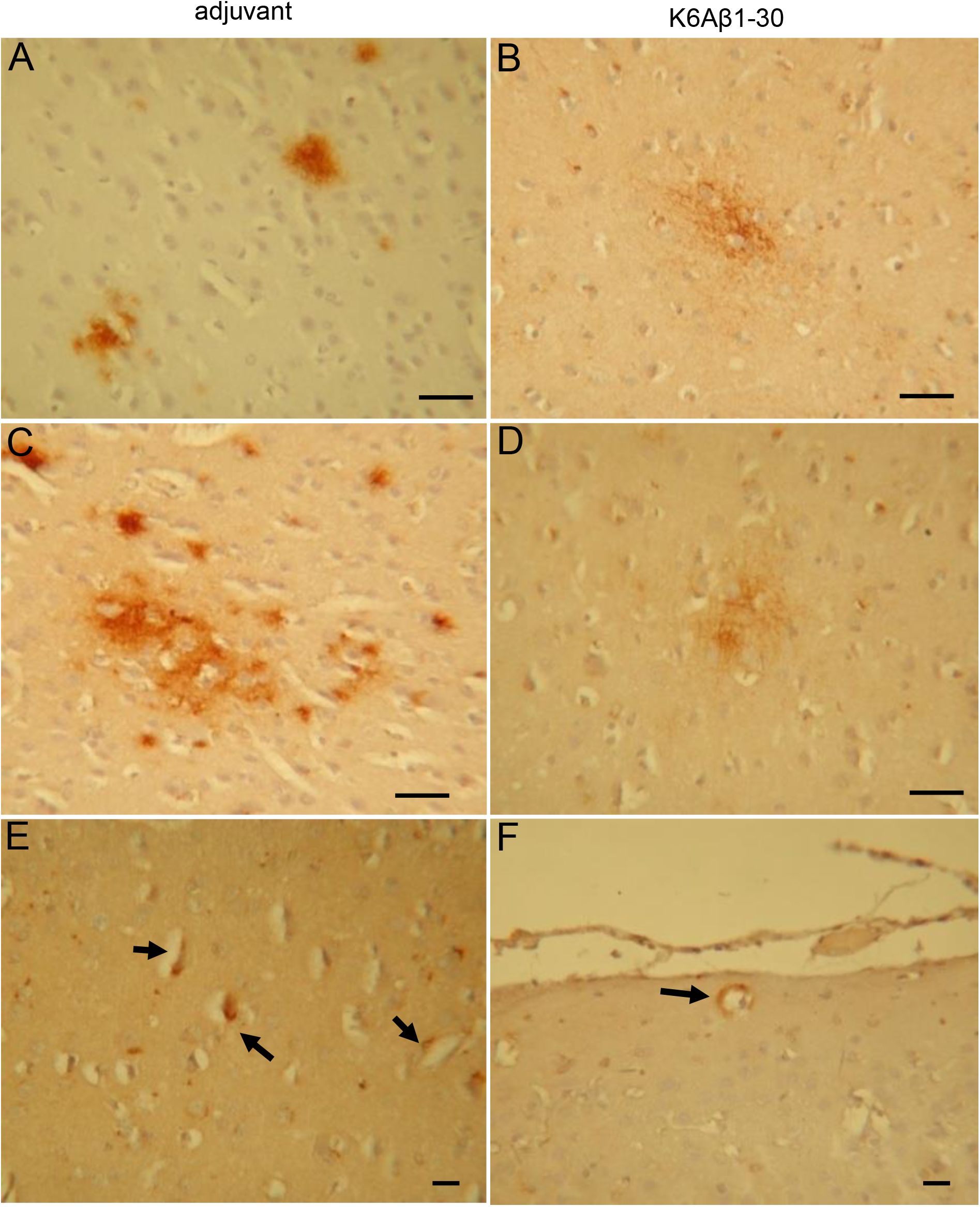
Amyloid Aβ deposits into the brain of mouse lemurs. Diffuse plaques stained with anti-Aβ1-42 (A, B) and with 4G8 antibodies (C, D). E, F: Aβ1-42 amyloid deposits into neurons and vessel walls (black arrows); bar : 30 µm.

Vascular Aβ deposits (accumulations in vessel walls) were detected in 38% of adjuvant-treated animals and 50% of immunised ones, and was most prominent in animals with extracellular amyloid deposits (Fig. 6E, F).

Intracellular Aβ was also detected in the brains of all mouse lemurs. It was most frequently seen in the soma of cortical pyramidal neurons (Fig. 7A, B) and sometimes in subcortical areas. No intracellular labelling was observed in the hippocampus. Interestingly, intracellular Aβ was decreased in several brain regions in immunised animals compared to adjuvant-treated group (Fig. 6C, occipital (p<0.05), frontal (p=0.06) and parietal cortex (p=0.108)).

**Figure 7.**
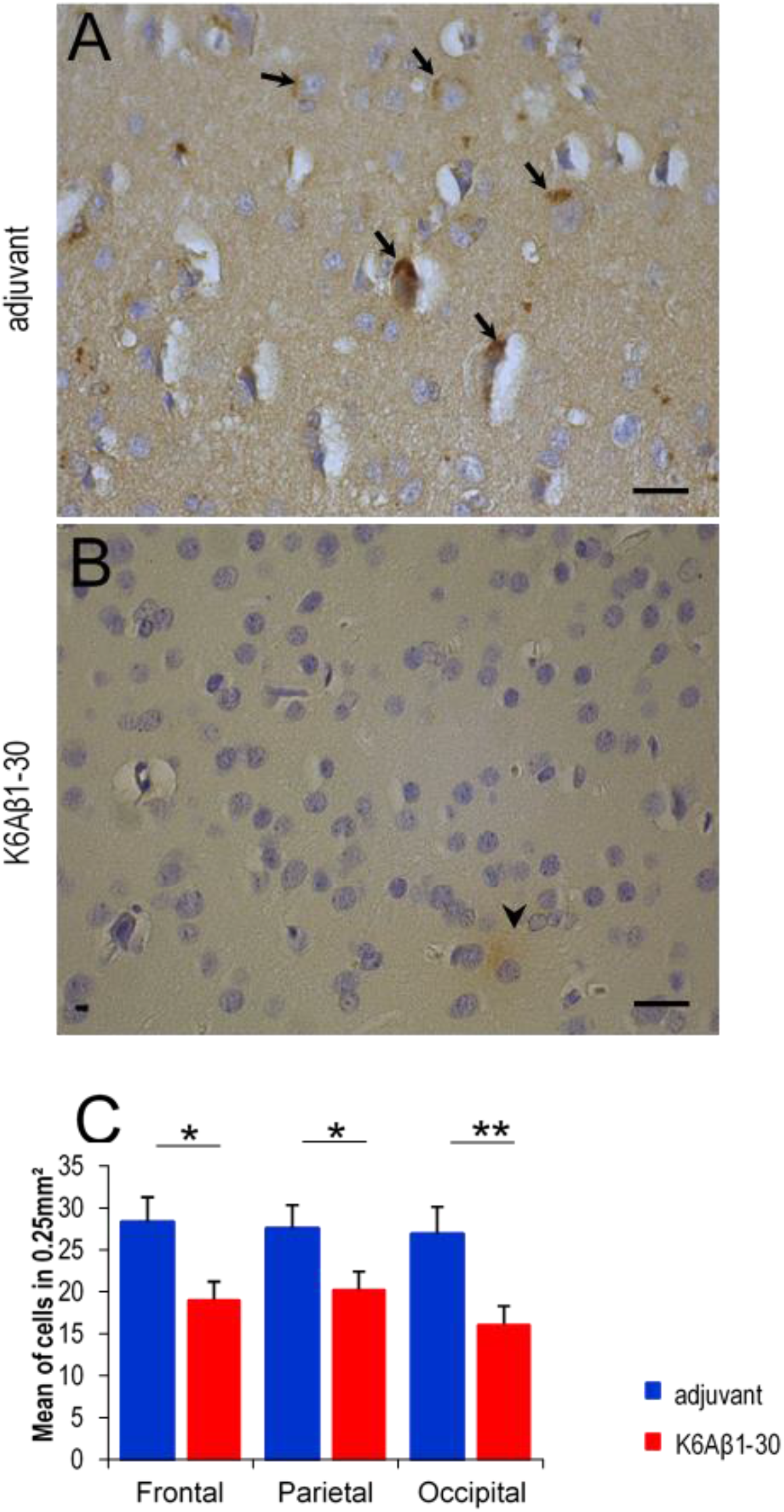
Intracellular Aβ in adjuvant-treated (A) and immunised (B) lemur brains. (A) numerous intracellular labelling (arrow) (B) absence of intracellular labelling ; only few external deposits (arrow head); C: Intracellular Aβ quantification in frontal, parietal, occipital cortex in control (n=16) and immunised lemurs (n=16). *p < 0.05 **p < 0.01 bar: 30 µm.

Taken together, these results suggest that the K6Aβ1−30 vaccination may have reduced Aβ burden in animals with such pathology.

### 3.4. K6Aβ1−30 immunisation had no impact on tau levels

The brains were analysed for tau and phospho-tau levels by Western blot and histological analysis to determine if the vaccine led to clearance of tau protein. Total tau was measured with monoclonal Tau-5 whereas phospho-tau was detected with monoclonal PHF1 and CP13 antibodies. The limited if any sarkosyl insoluble pellet of the brain homogenate indicates that the lemurs did not have much of insoluble tau. The soluble fraction analysis did not show any significant differences between adjuvant-treated and immunised mouse lemurs (Fig. 8). Specifically, analysis of the brain homogenate supernatant fraction showed no significant differences in phospho-tau (PHF-1, p = 0.43, CP13, p = 0.36), total tau (Tau-5, p = 0.14) or ratios (PHF-1/Tau-5, p = 0.41 and CP13/Tau-5, p = 0.73), between the two groups.

**Figure 8.**
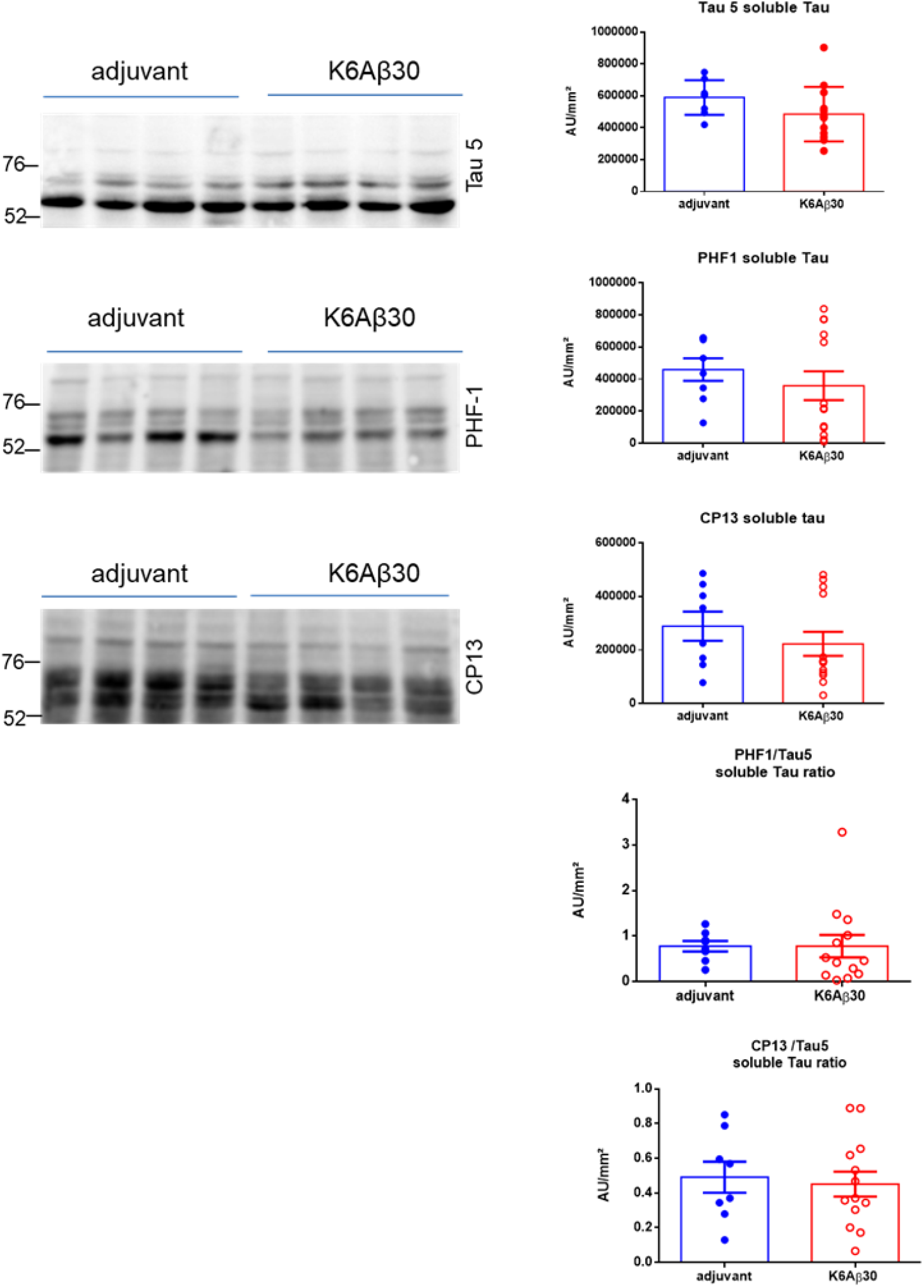
Tau levels on mouse lemur brains. Immunisation had no effect on soluble phospho-tau (PHF-1, CP13) total tau (Tau-5), or ratios on western blots of mouse lemur brain homogenates compared to adjuvant treated animals. Representative blots are shown on the left. Each bar represents the group average +/− SEM.

Immunohistochemical staining with CP13, PHF1 and AT100 antibodies revealed a mild tau pathology in mouse lemurs. CP13 and PHF1 immunoreactivity was more pronounced than AT100 staining. Tau staining was only observed in the two animals with Aβ deposits (one adjuvant treated and one immunised). The CP13 antibody revealed some pretangles in the hippocampus of the immunised and adjuvant-treated animals (Fig. 9A, B respectively). In the adjuvant treated animal, PHF1 stained neurons in the piriform cortex (Fig. 9C) and rare neurons were detected in the cortex with AT100 antibody (Fig. 9D). In the immunised animals no PHF1 or abnormally phosphorylated Tau labelling were observed.

**Figure 9:**
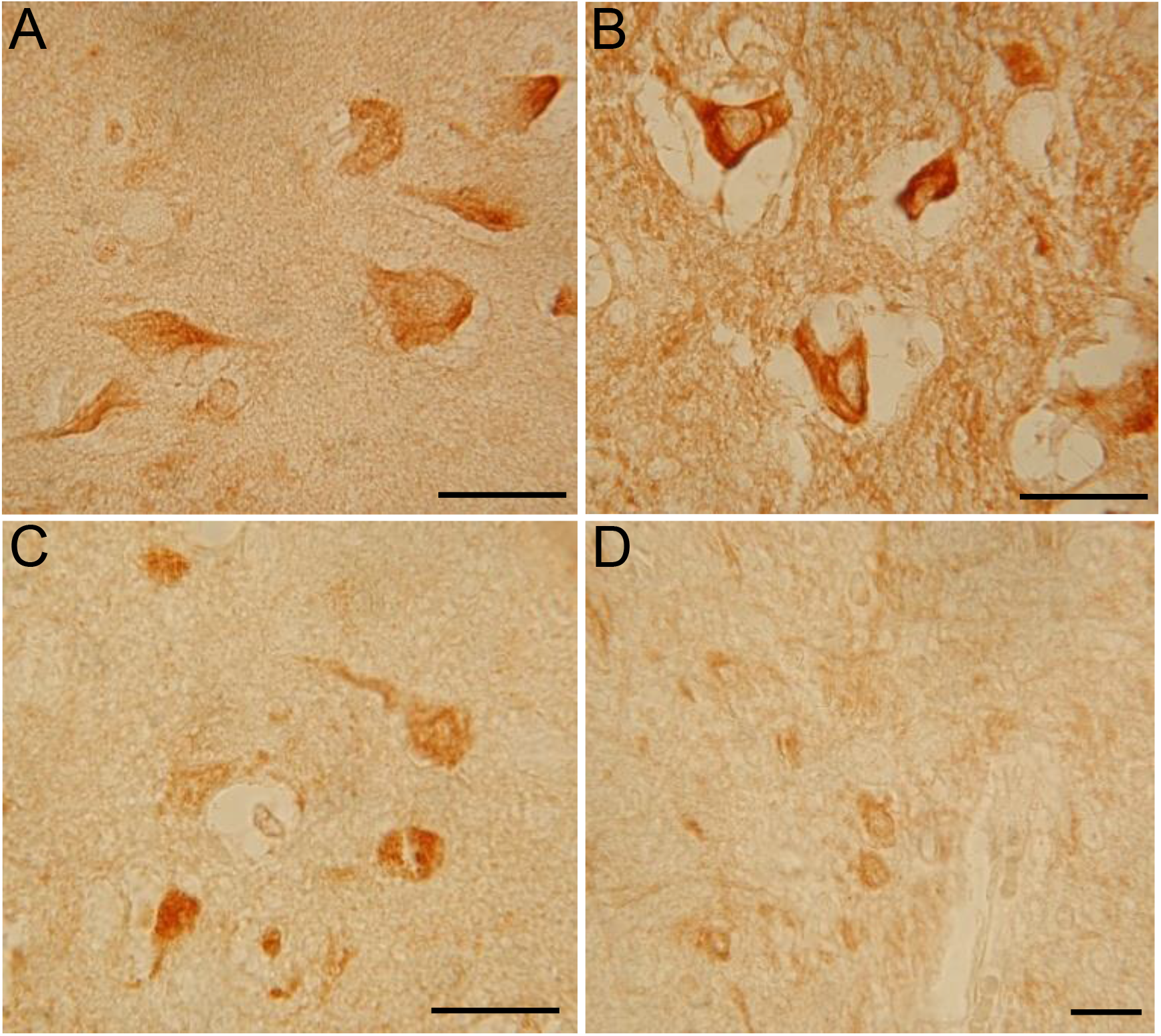
Tau labelling in mouse lemur brain. A-B: CP13+ neurons in hippocampus of immunised (A) or adjuvant treated (B) animals with Aβ plaques. Pathological tau in cortical neurons of the adjuvant treated animal evidenced by PHF1 (C) or AT100 (D) antibodies. Bar: 30 µm

The limited presence of pathological tau proteins in the brains of the mouse lemurs fits with the lack of insoluble tau in the brain homogenate following sarkosyl extraction, and did not allow us to conclude on the potential effect of the vaccine on tau pathology on tissue sections.

### 3.5. K6Aβ1-30 immunisation decreased microgliosis

We tried to quantify IL-4 and INFγ production by splenocytes after stimulation by the K6Aβ1-30 immunogen using a multi-analyte ELISArray kit. However, the OD values were below the detection levels in both treated and control animals (data not shown).

To address whether the vaccine treatment affected microgliosis and astrogliosis in the brains of mouse lemurs, we stained microglia and astrocytes with antibodies against ferritin and GFAP, respectively. We quantified the number of microglial cells in various cortical areas including frontal, parietal, occipital cortex and hippocampus. Classic antibodies used in rodents and humans to stain microglia such as Iba1, and lectin did not work under various conditions in Microcebus murinus. This is a well know issue in these animals. In Alzheimer human cortex, an antibody against ferritin has been shown to stain activated microglia [Kaneko et al, 1989]. We detected ferritin positive cells in cortical areas with a density higher in adjuvant-treated animals compared to immunised animals (Fig. 10 A, B). Double labelling Aβ1-42/ ferritin showed that microglia were associated with Aβ plaques (Fig. 10 C, D). The K6Aβ1-30 vaccine strongly reduced cortical and hippocampal ferritin positive cells by 46% (Fig. 10E, p < 0.01).

**Figure 10:**
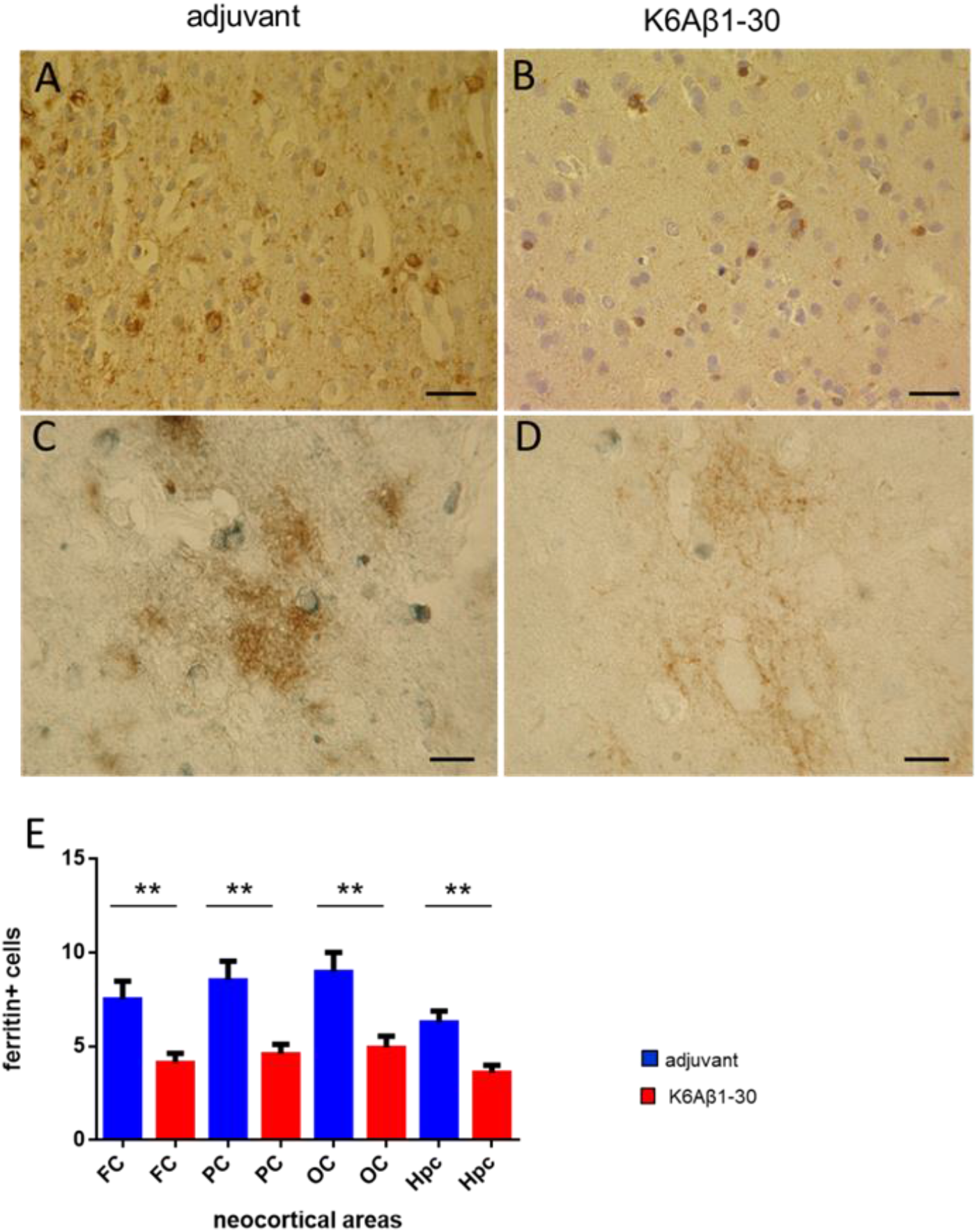
Staining of microglial cells with anti-ferritin antibody. A-B: Ferritin staining in animals with Aβ plaques. C-D : Double staining Aβ1-42 (in brown)/ ferritin (in black). A, C : Adjuvant-treated lemur with amyloid plaques. B, D: Immunised lemur with amyloid plaques. Bar: 30 µm. E: Ferritin quantification in adjuvant-treated (n=16) and immunised (n=16) lemurs. Immunised animals had less ferritin staining in frontal (FC), parietal (PC) occipital cortex (OC) and hippocampus (Hpc), compared to adjuvant treated lemurs (**p < 0.01).

The evaluation of astrocytes by GFAP antibody showed no difference between the treatment groups or between animals with (Fig. 11A, B) or without plaques (Fig. 11C, D) away from the plaques. As expected double labelling of GFAP and Aβ42 confirmed some astrogliosis in the vicinity of amyloid plaques in adjuvant and immunised animals respectively (Fig. 11 E, F).

**Figure 11:**
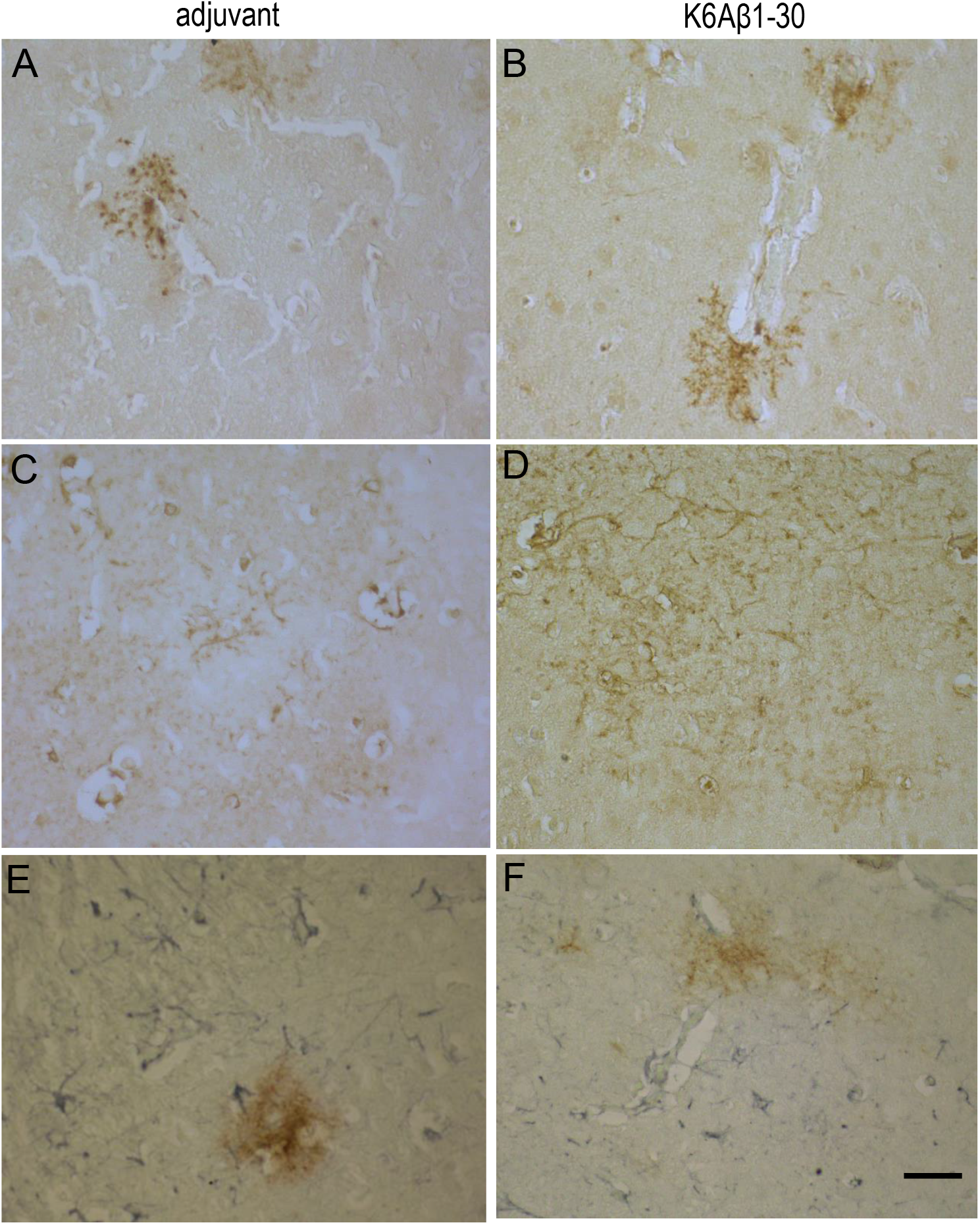
Astrogliosis. GFAP labelling in mouse lemur brain with (A, B) or without Aβ plaques (C,D). Double staining GFAP (black)/Aβ1-42 (brown) showing astrocytes in the vicinity of Aβ plaques. Bar: 30µm

Perl’s iron staining did not reveal any microhaemorrhages in either group. CD3 and CD45 immunostaining did not show accumulation of T cell in the brain of immunised or adjuvant treated mouse lemurs.

These results suggest that the immunisation strongly decreased microgliosis and that it was not associated with side effects that could have been revealed by the other markers.

## 4. Discussion

On the basis of the amyloid hypothesis, removal of Aβ aggregates by immunotherapy has been suggested as a treatment option to slow or halt AD ^35^. Immunotherapy focuses on the generation (active) or use (passive) of antibodies targeting a specific antigen, Aβ in this context, counteracting the disease by antibody-mediated removal of Aβ. An advantage of active immunisation is that a few vaccinations allow the patient to produce a prolonged antibody response. The variability of the induced response across patients is, on the other hand, a problematic aspect, especially when it comes to the elderly. In addition, undesirable side effects may occur after active immunisation, in particular when a T-cell response is induced, which can result in inflammation. Furthermore, with age, the competence of the immune system decreases and the likelihood of developing autoimmune responses is increased. Various Aβ immunotherapy strategies have been investigated for their therapeutic impact in preclinical rodent models of AD ^35–39^. Many of these have demonstrated beneficial effects in preventing, halting or reversing AD pathologies and preserving memory and learning functions. These promising effects were typically reported in pre and early-symptomatic AD transgenic mice, and have rarely been investigated in old late-stage mouse models for AD.

Interestingly, our prior studies in transgenic Tg2576 Aβ plaque mice using this same immunogen and adjuvant, K6Aβ1-30 in alum adjuvant, showed a robust antibody response, Aβ clearance and cognitive benefits in mice treated from 12-19 or 11-24 months ^9,27^, but a complete lack of effect in all of these parameters in mice treated from 19-24 months ^27^;.This particular immunogen has also led to robust Aβ clearance in that Tg2576 mouse model when administered as an oral vaccine expressed in attenuated Salmonella, with treatment lasting from 3-24 months ^40^. Aβ deposition begins in this mouse model around 12 months of age and is well established at 19 months. These findings if translatable to humans suggests that a stronger adjuvant and/or a multimer of the immunogen may be needed if treatment is initiated in old individuals. The relatively modest antibody response towards Aβ1-40 in the primates in the present study, without apparent side effects, suggests that they could have benefitted from such modification of the approach.

Aging is the major risk factor for AD development 24 and spontaneous models of AD offer the potential to bridge the translational gulf between promising rodent studies and failed human clinical trials ^38^. Non-human primate models have played a vital role in aging research as they manifest many of the structural and physiological modifications in the brain linked to chronological aging. Aβ immunisation has been tested in a limited number of non-human primate studies ^12,18,41,42^ and rarely on elderly monkeys ^17,33,43–45^. These reports have primarily focused on the production and characterization of Aβ targeting antibodies and their effect on cerebral Aβ clearance. Only one of these studies investigated the effect of the vaccine on cognition because of the difficulty of having access to aged monkeys 45. Since old mouse lemurs are easier to work with than aged larger primates, these animals may provide greater insights into the potential translation of efficient therapies, mimicking the individual variability found in humans.

Our present findings in a relatively small group of old primates indicate that active immunisation with an Aβ derivative (K6Aβ1−30-NH2) appears to be safe as a prophylactic measure. The vaccine did not appear to have detrimental effects on the general health of these old animals. Some animals died or had to be euthanized during the protocol in both groups (5 controls and 4 immunised, see Table 1). These deaths primarily occurred in the oldest mouse lemurs, which are known to suffer from a syndrome that leads to renal insufficiency, loss of weight and eventually death ^46^. At the autopsy, we observed nephritis in both groups, which is very common in untreated old mouse lemurs. Therefore, this phenomenon is unlikely to have been related to the adjuvant or the immunogen.

Initially, we had planned to examine IgG and IgM response to K6Aβ1-30, Aβ1-40 and Aβ1-42. However, the small amount of blood that could be retrieved from the animals without causing overt stress did not allow us to measure antibody binding to both Aβ1-40 and Aβ1-42. The main B-cell/antibody epitopes within the immunogen, K6Aβ1-30-NH2, Aβ residues 1-11 and 22-28, are found within both Aβ1-40 and Aβ1-42. Therefore, antibody binding to Aβ1-40, compared to the immunogen itself, provides sufficient information on antibody response to any Aβ peptide that contains its first 30 amino acids, as is the case for both Aβ1-40 and Aβ1-42. Although the immunisation elicited a relatively low antibody titre against Aβ1-40, the mouse lemurs did have a rather robust IgG response that decreased after 6 months. Re-immunisation induced high level of antibodies in half of the animals, suggesting than an optimal protocol for maintaining an antibody response would be repeated immunisations with no longer than 6 month intervals. Kofler and colleagues ^43^ have found that, in immunised aged responder macaques, the antibody responses were significantly lower than in juvenile animals. Compared to our previous report on young mouse lemurs, our old animals had a better IgG and IgM response after re-immunisation than the young animals ^12^. Relevant to vaccines is the observation that, regardless of age, females tend to show greater antibody responses than males ^47^. However, in this group of old mouse lemurs, the sex of the animals could not explain the difference between responders and non-responders.

Early detection of age-associated cognitive dysfunction is crucial, as this provides a window of opportunity to understand the breakdown of brain systems and to implement interventions that may limit the progression of disease. The difference in cognitive performances observed before and after treatment could provide insights on vaccination efficiency. Hara and colleagues ^45^, with a finger maze test in aged cynomolgus monkey were not able to see effect of their AAV-Aβ vaccine. In the present study, a trend for improvement in spatial memory was observed in the old mouse lemurs during immunisation, pointing to functional benefits of the vaccine, probably related to Aβ clearance. We have shown recently, using pairwise discrimination and reversal learning test in old mouse lemurs that impaired discrimination learning is linked to extracellular cortical accumulation of Aβ and that subjects with intraneuronal Aβ aggregates fail more often in a pre-training task ^48^. Although the antibody response towards Aβ was relatively modest, the immunisation appeared to result in a reduction of the extracellular Aβ burden, similar to what was observed in the neuropathological analyses from AN1792 recipients that showed a lower mean Aβ load compared to an age-matched non-immunised controls ^49^. Intracellular Aβ was significantly decreased in the immunised lemurs and was half that of the adjuvant treated animals, and even less when compared with the 7 age-matched controls. The intracellular deposition of Aβ prior to its extracellular accumulation has been described in the brains of both AD patients and animal models ^50^, and a correlation between intracellular Aβ and neurodegeneration was recently reported in a triple-transgenic AD (3xTg-AD) murine model ^51^. Furthermore, a decrease of intraneuronal Aβ after Aβ targeting vaccination has been observed in transgenic mice ^51^ and in macaque brain ^45^. The lack of correlation between the antibody response and plasma or brain levels of Aβ suggests that the study was not sufficiently powered to detect such association.

One of the main challenges of immunotherapy is understanding the precise role of neuroinflammation. The measurement of pro-inflammatory cytokines were below the detection levels in both treated and control animals, and the proportion of GFAP-immunoreactive areas and signal intensities did not show any difference between groups. However, the proportion of ferritin-positive cells (principally microglia) in hippocampus and neocortex was reduced by half in the vaccinated lemurs compared to the controls that received adjuvant alone. This appears to bode well for the safety of this immunogen.

## 5. Conclusions

This study in aged lemur primates indicated that the K6Aβ1-30 vaccine was safe. It showed a trend to improve spatial memory in the animals that was associated with reduced microgliosis, and clearance of intracellular Aβ from the brain. These observations offer a compelling and testable hypotheses that could form the basis of clinical studies.

## Acknowledgments

The authors want to thank the late Peter Davies for providing tau antibodies. The authors further like to thank the breeding facility from the University of Montpellier (RAM-CECEMA) as well as the animal keepers, Sylvie Rouland, Joel Cuoq, Faustine Hugon and Méline Péguet.

This research was funded by grant from the National Institutes of Health (R01 AG020197) and by MMDN.

