## supplementary figures for "Amyloid-β targeting immunisation in aged non-human primate (*Microcebus murinus*)"

Supplementary data

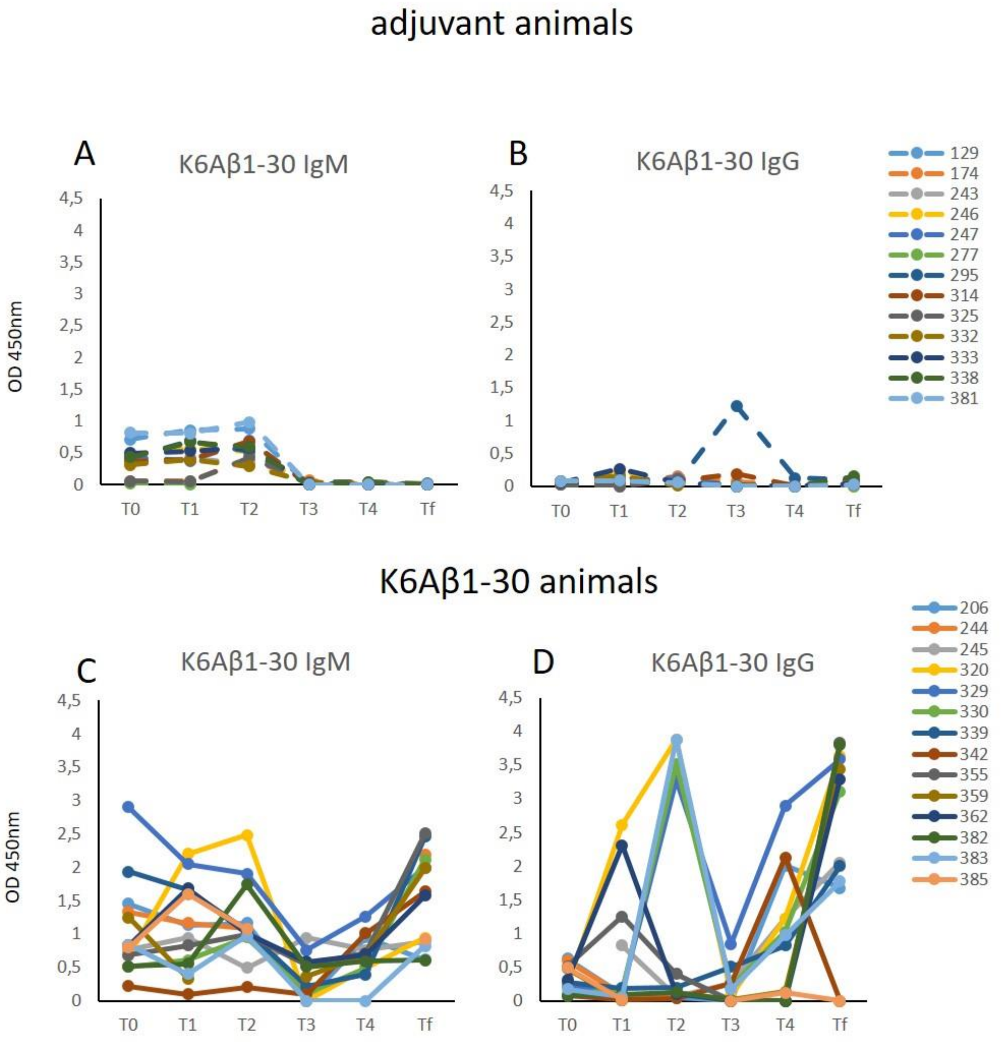


***Figure 1 Suppl: Individual antibody response in adjuvant treated** (A, B) **and K6Aβ1-30 treated** (C, D) **mouse lemurs**. **K6Aβ1-30 IgM (A,C) and K6Aβ1-30 IgG (B,D)** The y-axis depicts the absorbance at 450 nm of antibodies bound to K6Aβ1-30 coated onto the plates. Animals with limited blood samples are not included (n=3 and n=2 for adjuvant- and K6Aβ1-30 treated animals, respectively). The legend shows the assigned animal numbers.*

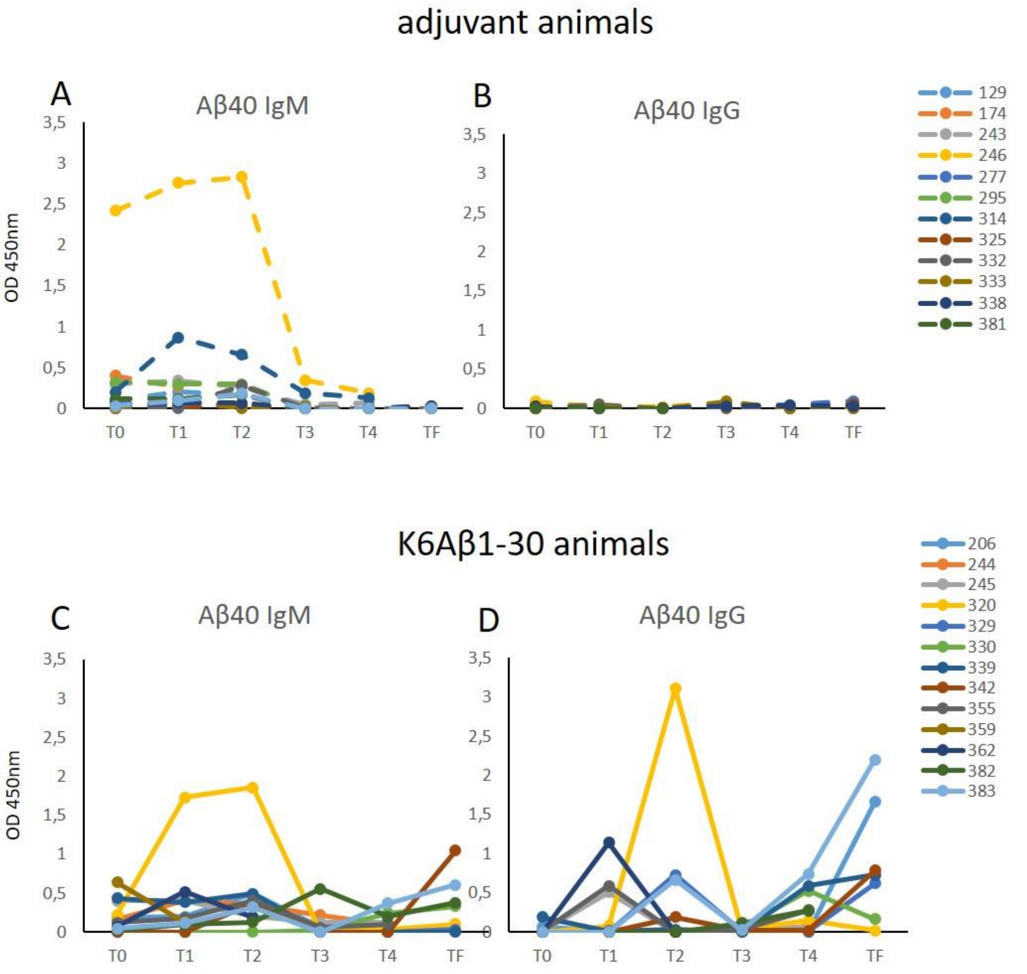


***Figure 2 Suppl: Individual antibody Response in adjuvant treated*** *(A, B)* ***and K6Aβ1-30 treated*** *(C, D)* ***mouse lemurs****.* ***Aβ40 IgM (A,C) and Aβ40 IgG (B,D)*** *The y-axis depicts the absorbance at 450 nm of antibodies bound to Aβ1-40 coated onto the plates. Animals with limited blood samples are not included (n=4 and n=3 for adjuvant- and K6Aβ1-30 treated animals, respectively). The legend shows the assigned animal numbers.*
